# Investigating temporal and spatial variation of eDNA in a nearshore rocky reef environment

**DOI:** 10.1101/2020.12.29.424660

**Authors:** Taylor Ely, Paul H. Barber, Zachary Gold

**Affiliations:** Department of Ecology and Evolutionary Biology, University of California Los Angeles, Los Angeles, California, USA; School of Life Sciences, University of Hawaii Manoa, Honolulu, Hawaii, USA

**Author notes:** Corresponding author, (TE).

## Abstract

Environmental DNA (eDNA) is increasingly used to measure biodiversity of marine ecosystems. However, key aspects of spatial and temporal dynamics of eDNA remain unknown. Particularly, it is unclear how long eDNA signals persist locally in dynamic marine environments, since degradation rates have predominantly been quantified through mesocosm studies. To determine *in situ* eDNA residence times, we introduced an eDNA signal from a non-native fish into a Southern California rocky reef ecosystem, and then measured changes in both introduced and background eDNA signals over 96 hours. Foreign eDNA signal could no longer be detected 7.5 hours after introduction, far exceeding disappearance rates quantified in laboratory studies. In addition, native vertebrate eDNA signals varied greatly over the 96 hours of observation, but time of day and tidal direction did not drive this variation in community structure. Species accumulation curves showed that standard sampling protocols using 3 replicate 1 L sea water samples were insufficient to capture full diversity of local marine vertebrates, capturing only 76% of all taxa. Despite this limitation, a single eDNA sample captured greater vertbrate diversity than 18 SCUBA based underwater visual transect surveys conducted at a nearby site. There was no significant difference in species richness between temporal replicates and spatial replicates, suggesting a space for time substitution may be effective for fully capturing the diversity of local marine vertebrate communities in nearshore rocky reef environments. This result is particularly important in designing eDNA metabarcoding sampling protocols to capture local marine species diversity.

## Introduction

Environmental DNA (eDNA) is increasingly used to investigate biodiversity of marine ecosystems [1]. eDNA is produced when organisms shed genetic material into the environment [2]; by isolating, extracting, and sequencing this eDNA, resident marine species can be identified through metabarcoding [1]. Recent studies demonstrate that eDNA techniques can outperform traditional visual census surveys in species detection [3,4], particularly for cryptic and rare species [5], while at the same time being noninvasive and cost effective [2]. As such, eDNA represents a promising alternative to traditional biodiversity surveys, which are time and labor intensive, require substantial taxonomic expertise, and can pose significant safety hazards to researchers [2,6].

Despite the promise of eDNA, much remains unknown about the dynamics of eDNA in the environment. eDNA degrades in the environment due to a combination of abiotic (e.g. temperature, UV, and pH) and biotic (e.g. microbial activity) processes [7]. Previous studies report that eDNA of marine fishes degrades in a laboratory setting on a scale of 0.5 to 7 days, but usually around 3-4 days [3,8]. In nature, however, water transport and water mixing affect the persistence and detection of eDNA in the water column, processes that are fundamentally altered in laboratory settings [8].

To date, most eDNA studies examining eDNA transport have focused on single species in freshwater systems characterized by relatively simple flow dynamics such as spawning salmon in streams [e.g. 9,10]. However, recent work investigating the spatial and temporal variation of *Pseudocaranx dentex* (white trevally) in Maizuru Bay, Japan found that eDNA signatures fell below detection thresholds under just 2 hours after the removal of a foreign eDNA source [11]. This result indicates that eDNA signatures in marine systems can fall below detection limits much faster than reported in laboratory experiments, suggesting that a combination of degradation, advection, generation, dispersion, and/or diffusion in marine systems dramatically differs from laboratory experiments. However, it is unclear whether this result is generalizable to all marine ecosystems, including temperate or polar ecosystems where colder water temperatures could slow degradation processes.

In dynamic aquatic ecosystems, the combination of eDNA degradation and transport can result in temporal variation in eDNA signatures [12]. As such, it is essential to understand temporal variation of eDNA in marine environments as well as the spatial scale of eDNA variation so that proper eDNA sampling strategies can be developed and results can be properly interpreted. Our present understanding of short-term temporal variation in eDNA signatures of entire marine vertebrate communities is limited to a single study on an intertidal ecosystem. Kelly et al. [13] found that tides did not have a strong or consistent effect on community composition, but that temperature and salinity did have a significant effect, suggesting that the movement of water masses—rather than tides alone—has the strongest effect on eDNA signatures.

Transport of eDNA in marine environments may not strongly impact local eDNA signatures if the eDNA only persists for a few hours, producing a highly localized signal. Such highly localized signals are reported in multiple studies. For example, Port et al. [14] found that marine vertebrate communities differed on a 60-100 m scale. This small spatial scale of community differences could be the result of either limited transport of eDNA due to the unique geographic and benthic topography of Lovers Cove, Pacific Grove, CA or due to the rapid degradation rates of marine eDNA. Recent work building off this study in more dynamic marine environments found that eDNA signatures of marine communities displayed spatial variation on the scale of hundreds of meters to a few kilometers [15]. Similarly, Yamamoto et al. [16] found significant spatial differences occurred around 800 m. Together these results suggest that eDNA has strong spatial variation in marine ecosystems.

Understanding the temporal and spatial variation of eDNA in marine ecosystems is becoming increasingly important, as eDNA is being viewed as a potential alternative to traditional visual survey methods used in the monitoring of coastal marine ecosystems [2,5]. A key potential advantage of eDNA is the ability to efficiently detect a larger number of taxa compared to visual surveys [3,4,14] as visual surveys frequently focus on a small subset of indicator taxa due to logistical difficulties [17]. For example, out of at least 178 fish species that inhabit Southern California kelp forest [18], Reef Check California only monitors 33 of these species in their visual surveys [19]. However, for eDNA to be maximally useful, it is essential to understand the temporal and spatial variation of eDNA to ensure proper sampling protocols to reliably detect these indicator species and the rest of the marine vertebrate community.

To better understand temporal and spatial variation in eDNA signatures, this study investigates the *in situ* persistence of eDNA in a nearshore rocky reef habitat using a Eularian sampling regime. First, we examine the persistence of a foreign eDNA signature after introduction in a given location. Second, we investigate how natural eDNA signatures at this location fluctuated over the duration of the study, and compare these results to visual survey protocols employed in Southern California waters by local kelp forest monitoring programs. Combined, this approach will provide critical insights into the dissipation of eDNA signatures over time and space, and how sampling can be optimized to account for this variation, providing the most robust eDNA data for monitoring marine vertebrate communities.

## Methods

### Sample collection and filtering

To create a foreign eDNA signature, we homogenized one raw filet (414 grams) of *Ctenopharyngodon idella* (grass carp) muscle tissue in 1L of MilliQ water (EMDMillipore, Burlington, MA) in a blender at high speed for 60 seconds. Tissue was used instead of PCR product because of recent evidence that eDNA derives from whole cells rather that freely associated DNA [8,20]. Due to the potential for seafood mislabeling [21], we DNA barcoded the sample to verify the species by including a tissue sample as a positive control.

We conducted fieldwork at the USC Wrigley Marine Science Center on Catalina Island, California, located in Big Fisherman’s Cove (33°26’42.43”N, 118°29’4.05”W). This field station sits within a protected bay and has a long dock to facilitate sampling along a fixed transect without disturbance from SCUBA divers. We established fixed sampling points along a 38 m transect. Location A was closest to shore, Location B was 19 m seaward, and Location C was 19 m further seaward. The depths of location A, B, and C were 7.3 m, 8.2 m, and 11.2 m respectively.

Prior to introducing the foreign eDNA signal, we established baseline eDNA signatures by collecting one-liter water samples at each site on SCUBA. SCUBA divers then released the total volume of homogenized *C. idella* tissue one meter above the sea floor at Location B on September 6^th^, 2017. We then collected one-liter water samples for eDNA analysis at each of the three sampling locations for a period of 96 hours. For the first 12 hours, we collected samples every 1.5 hours. After this initial period, we sampled only at 24, 48, and 96 hours after the release of the foreign eDNA. Sterile protocols were followed throughout following the guidelines of Goldberg et al. [22].

To eliminate diver related introduction of *C. idella* during sample collections, all samples following the introduction of the foreign eDNA signature were taken from the dock using a 4-liter Niskin bottle hand-lowered to 1 m above the sea floor. At the surface, we transferred 1 liter of seawater from the Niskin bottle into a sterile 1000 mL kangaroo gravity feeding bag (Covidien, Dublin, Ireland, Product Number 8884702500). This bag was immediately placed in a cooler on ice packs (−20°C) and transported to the lab, less than 300 m away, for filtration.

To filter eDNA from the water samples, we fitted sterile 0.635 cm diameter Nalgene tubing with a luer-lock adapter to the pouches and connected the sample to a 0.22 μm diameter PVDF Sterivex filter unit (EMDMillipore, SVGPL10RC) [23]. We then hung the bags and attached filters in the lab, allowing samples to gravity filter for a maximum of 40 minutes (S1Table). Following eDNA filtration, we stored filters at −20°C and transported the filters to UCLA for molecular laboratory work.

To ensure that no DNA carried over between sampling events, we cleaned the Niskin bottle between sample collections by rinsing the bottle with surface water above each location for 30 seconds [24]. To test for contamination, we ran a field blank which consisted of 1 L of nuclease-free water placed inside the Niskin bottle previously rinsed with locally sourced tap water for 30 seconds. This water was then transferred to the gravity feeding bag for filtration and processed identically to field samples.

### DNA extraction

We extracted DNA from the Sterivex filters using a modified Qiagen DNeasy Blood and Tissue Kit (Qiagen Inc., Germantown, MD) following protocols from Spens et al. [25]. The modifications include the following steps. We added 80 μL of proteinase K and 720 μL ATL buffer from the kit directly into the Sterivex filter before sealing both ends of the filter. We then placed the filters in a rotating incubator overnight at 56°C. Following incubation, we removed the liquid from each Sterivex filter using a sterile 3 mL syringe and transferred the solution into 1.5 mL tubes. We then added equal parts AL buffer and Ü°C ethanol to an equal volume of extracted liquid. The eDNA sample was then extracted with the Qiagen DNeasy Blood and Tissue Kit without any further modifications to the manufacturer’s protocol.

### PCR amplification and DNA sequencing

We amplified the *12S* region of mitochondrial DNA using MiFish Universal Teleost specific primers modified with Illumina Nextera adapter sequences (MiFish-U, S2 Table) [23,26]. This primer set primarily targets teleost fish but also can detect a wide variety of other vertebrates, including marine mammals and birds [26]. PCR reaction volume was 25 μL and included 12.5 μL Qiagen 2x Multiplex Master Mix, 2.5 μL MiFish-U-F (2 mM), 2.5 μL MiFish-U-R (2 mM), 6.5 μL nuclease-free water, and 1 μL DNA extraction. PCR thermocycling employed a touchdown profile with an initial denaturation at 95°C for 15 min to activate the DNA polymerase followed by 13 cycles with a denaturation step at 94°C for 30 sec, an annealing step with temperature starting at 69.5°C for 30 sec (temperature was decreased by 1.5°C every cycle until 50°C was reached), and an extension step at 72°C for 1 min (S3 Table). Thirty-five additional cycles were then carried out at an annealing temperature of 50°C using the same denaturation and extension steps above, followed by a final extension at 72°C for 10 min (S3 Table). All PCR experiments included negative controls. We then confirmed successful PCR amplification through gel electrophoresis on 2% agarose gels.

To prepare the sequencing library, we first pooled 5 μL from each of the 3 PCR technical replicates. We then purified these PCR products, removing strands less than 100 bp long, using Sera-Mag (Sigma-Aldrich) bead protocol and eluted the purified product in 40 μL nuclease-free water [27]. We attached Illumina Nextera indexing primers to purified products through an indexing PCR [28]. The PCR reaction volume per sample was 25 μL and comprised of 12.5 μL Kapa HiFi Hot Start Ready Mix, 0.625 μL Primer i7, 0.625 μL Primer i5, 6.25 μL nuclease-free water, and 5 μL template (~5 ng). The thermal cycle profile started with 95°C for 5 minutes followed by 5 cycles of denaturing at 98°C for 20 seconds, annealing at 56°C for 30 seconds, and extension at 72°C for 3 minutes. Thermal cycling concluded with a final extension of 72°C for 5 minutes. These indexed samples were again purified to remove strands less than 100 bp long using Sera-Mag beads as described above. DNA concentrations were quantified using the BR Assay Kit (Thermofisher Scientific, Waltham, MA, USA) on a Victor3 plate reader (Perkin Elmer, Waltham, MA, USA). We then generated the final library by pooling equal concentrations of DNA from all indexed samples. The library was then sequenced at UC Berkeley’s QB3 Genomics in a single Illumina MiSeq Paired end 300×2 sequencing run.

### Bioinformatics

We analyzed the resulting sequences using the *Anacapa Toolkit* (version: 1) [23] identifying the number of reads of *C. idella* and amplicon sequence variants (ASVs) from the native vertebrate communities. We used the standard *Anacapa Toolkit* parameters with the *CRUX* generated *12S* reference library as described in Curd et al. [23] with the addition of 757 barcodes of California fish species [29]. Taxonomic assignment was determined with a Bayesian confidence cutoff score of 60 [30].

We employed an established decontamination package, *Gruinard Decon* (version 0.0), using *microDecon* (version 1.0.2) [31,32], in R (version 3.6.1) [33] and followed index hopping removal using the methods Kelly et al. [13]. We then normalized our data using the eDNA index metric following the methods Kelly et al. [34]. This metric assumes that PCR biases originate from template-primer interactions which remain constant across eDNA samples and thus allow us to infer relative abundance changes of a single taxa between samples [34].

### Statistical analysis

To examine degradation of the introduced eDNA, we plotted the index of *C. idella* reads at each time point across the first 24 hours of sampling using R (version 3.6.1) [33]. Unlike laboratory studies, we chose not to fit an exponential model to the data as the *C. idella* reads did not decrease in a consistent pattern [3,8]. For comparisons of native vertebrate communities over time, we first visualized which taxa were present in each location (A/B/C, N=3) and time point (0-96 hr, N=12) by generating a heat map using the R package *phyloseq* (version 1.28.0) [35]. We generated two separate heat maps, one using all taxa detected and a second using only a subset of species monitored by one or more of local kelp forest monitoring programs (National Park Service Kelp Forest Monitoring Program (KFM), Partnership for Interdisciplinary Studies of Coastal Oceans (PISCO), and Reef Check) (S4 Table).

To test the underlying factors shaping temporal variation in native eDNA community signatures, we investigated how species communities changed in response to three variables: direction of tide (incoming/outgoing/peak, N=3), location (A/B/C, N=3), and time point (0-96 hr, N=12). We conducted a PERMANOVA test on the Bray-Curtis dissimilarities calculated between each sample to determine the effect of each variable on community composition using the R package *vegan* (version 2.5-6) [36]. We chose Bray-Curtis dissimilarities over Jaccard dissimilarities following the methods of Kelly et al. [34] given that eDNA index enables us to infer changes in relative abundance between taxa.

To compare species richness recovered with spatial versus temporal replicates, we calculated the means and 95% confidence intervals (CI’s) using the R package *iNext* (version 2.0.20) [37] for sets of three random samples. *iNext* determines sets of spatial replicates from mean taxa recovered at a given time point across all three sampled locations and sets of temporal replicates from mean taxa recovered at any given location across 3 randomly selected time points. Mean and 95% CI calculations were repeated for the subset of monitored species detected.

Lastly, to determine the number of eDNA samples needed to capture subsets of species diversity, we calculated species accumulation curves using R package *iNext* (version 2.0.20) [37]. We did this to determine how many samples are required to recover 1) all species detected by eDNA, 2) a subset of species monitored by local kelp forest monitoring agencies (KFM, PISCO, and Reef Check), and 3) a subset of species only monitored by Reef Check (S4 Table). We focus specifically on Reef Check because they monitor a site ~100 m away from our site at the Wrigley Marine Science Center, providing a comparison between eDNA results and traditional visual survey results (S4 Table). We created all graphs and performed all calculations using R (version 3.6.1) [33] with *phyloseq* (version 1.28.0) [35], *Ranacapa* (version 0.1.0) [38], *treemapify* (version 2.5.3) [39], *iNext* (version 2.0.20) [37] and *vegan* (version 2.5-6) [36] packages.

## Results

### Sequencing

We generated 6,613,832 reads across 36 samples and 5 controls. The number of reads per sample ranged from 54,988-558,298 reads (average = 179,719 reads/sample), excluding controls. After decontamination steps, 754 ASVs were recovered.

### Temporal and spatial variation in foreign eDNA signatures

Results showed no *C. idella* eDNA in any samples prior to introduction of the tissue homogenate, but it was detected at all three sites at similarly low detection levels (eDNA index scores = 0.025-0.130) 1.5 hr after release. The strongest eDNA signature was detected 3 hr after release, but only at site C (eDNA index score = 1). eDNA index scores decreased over time in an inconsistent fashion. No foreign eDNA was detected at site A and C at 4.5 hrs, but it was detected at all sites at 6 hrs, with site C at 6 hrs having the second highest eDNA index score over the entire experiment (eDNA index score = 0.715). By 7.5 hrs, foreign eDNA was no longer detected (Fig 1).

**Fig 1.**
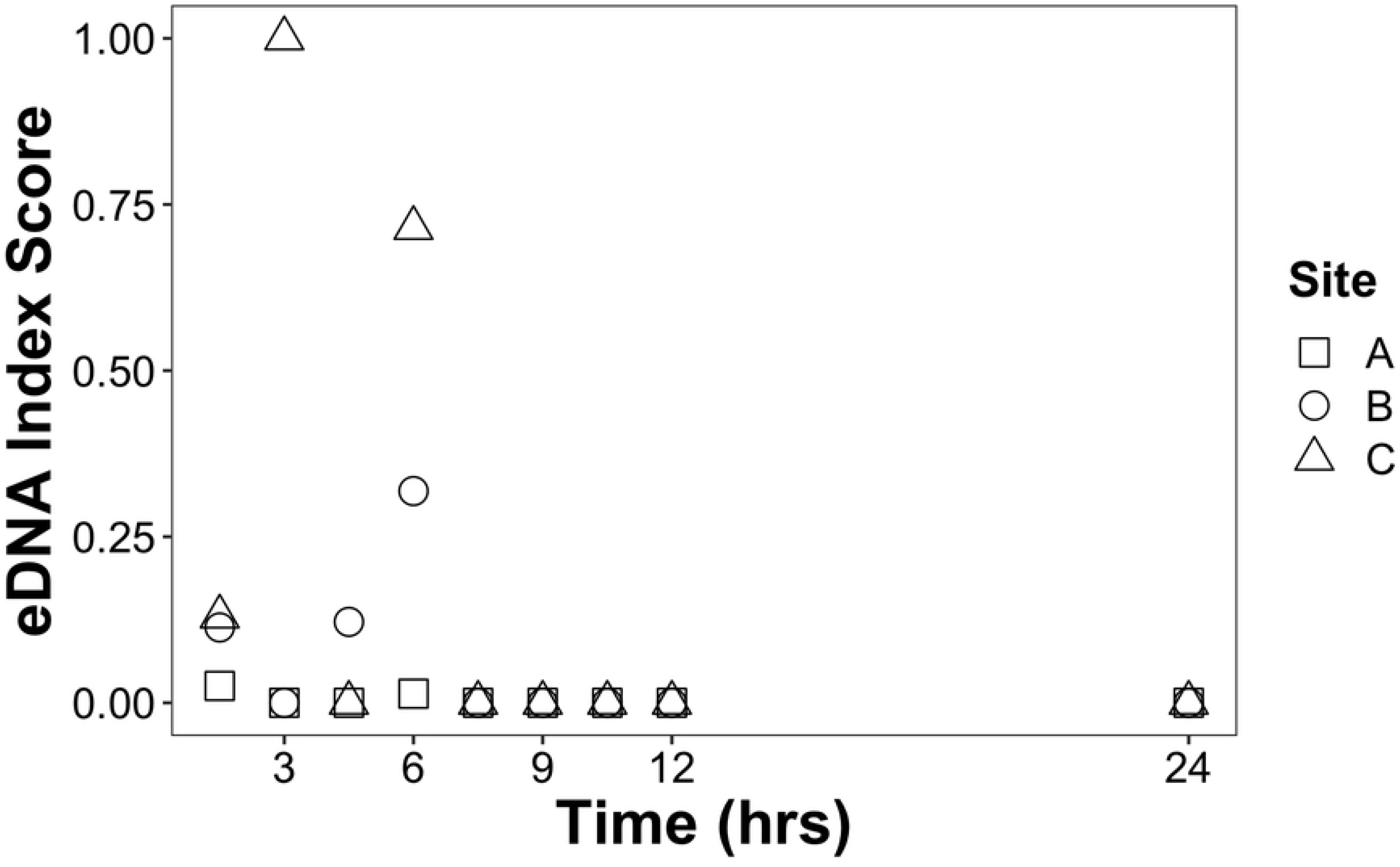
Detection of Foreign eDNA over Time. Plot of eDNA index for *C. idella* detected at locations A (squares), B (circles), and C (triangles) over time per location for the first 24 hours

### Temporal and spatial variation in native eDNA signatures

A total of 99 taxa were present in at least one sample across the 3 sample locations and 12 sampling times, spanning 4 classes, 29 orders, 57 families, 84 genera, and 85 species (S4 Table); however, only one species, *Chromis punctipinnis*, was detected in every sample (S1 Fig). The remaining taxa detected exhibited heterogeneous patterns and were absent from one or more sampling points and times; this pattern was also observed for the subset of species monitored by the KFM, PISCO, and Reef Check (Figs 2 and S2). We note that the presence of a spike in foreign eDNA did not reduce the detection of native taxa. The mean number of taxa detected when the foreign eDNA signature was present was 21 species (σ=6) and the mean number of taxa detected when the foreign eDNA signature was absent was 20 species (σ=5).

**Fig 2.**
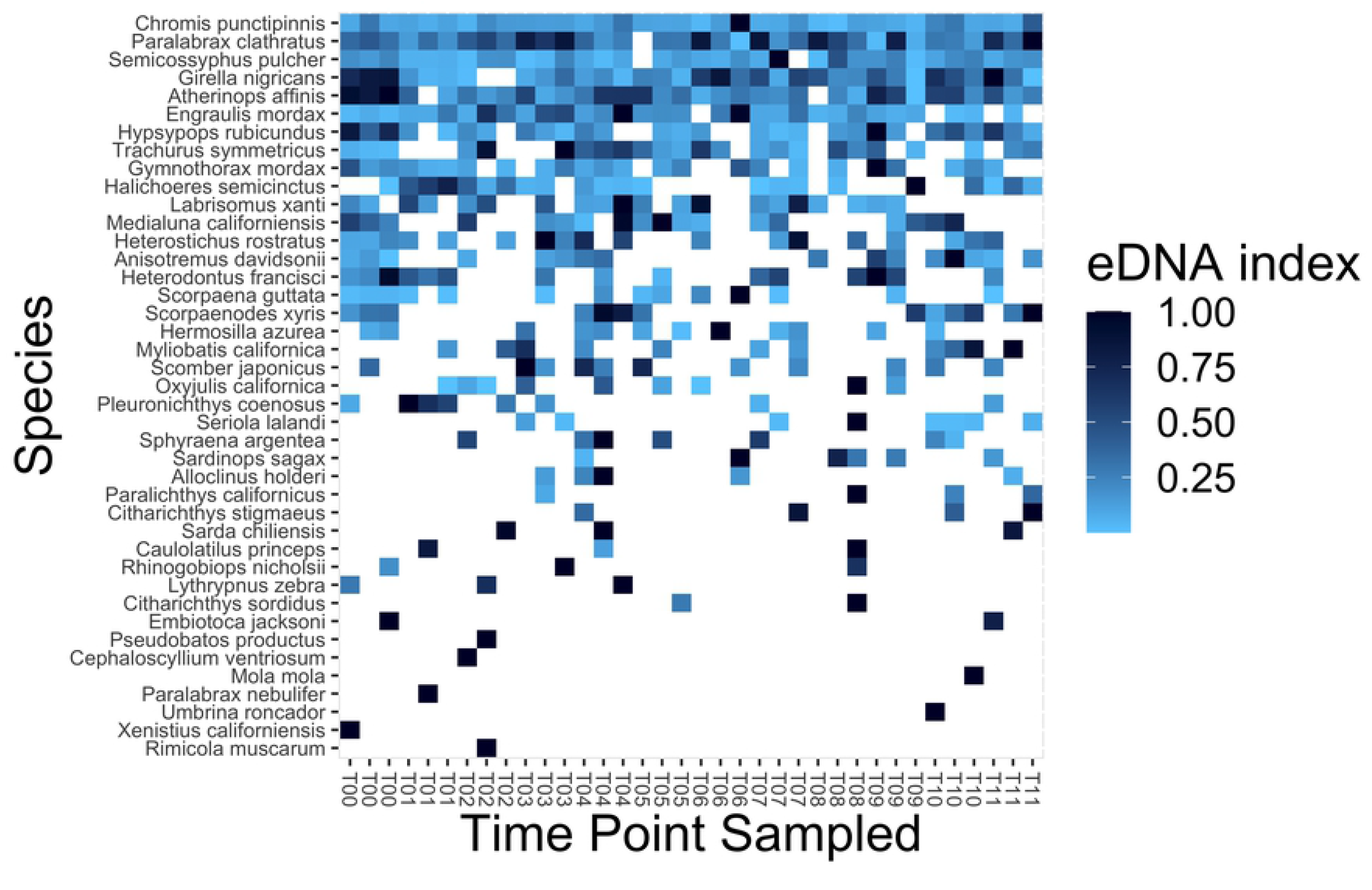
Heat Map of Species Monitored by KFM, PISCO, and Reef Check Detected Over Time. Heat map showing strength of the eDNA index for species monitored by KFM, PISCO, and Reef Check observed. Darker blue indicates higher index values. White indicates the species was not detected. Time point sampled is ordered by time and faceted by location (in order of location A, B, then C). Species are ordered by decreasing total detections.

Results from PERMANOVA found that time point accounted for the largest portion of variation in vertebrate assemblages (PERMANOVA; R^2^=32%) (Fig 3). The next most important sources of variation were direction of tide (PERMANOVA; R^2^=8%) and location (PERMANOVA; R^2^=7%) (Fig 3). The remaining 54% of variation was unaccounted for (Fig 3). Time point, direction of tide, and location of samples were all significantly correlated with Bray-Curtis community structure (PERMANOVA; F_9_=1.4634, F_2_=1.6107, and F_2_= 1.4482 respectively; p = 0.001, 0.002, and 0.005 respectively) (Fig 3). However, an NMDS ordination plot shows very weak clustering with no discernible effect of tides or time point (Stress > 0.25; S3A and S3B Figs respectively).

**Fig 3.**
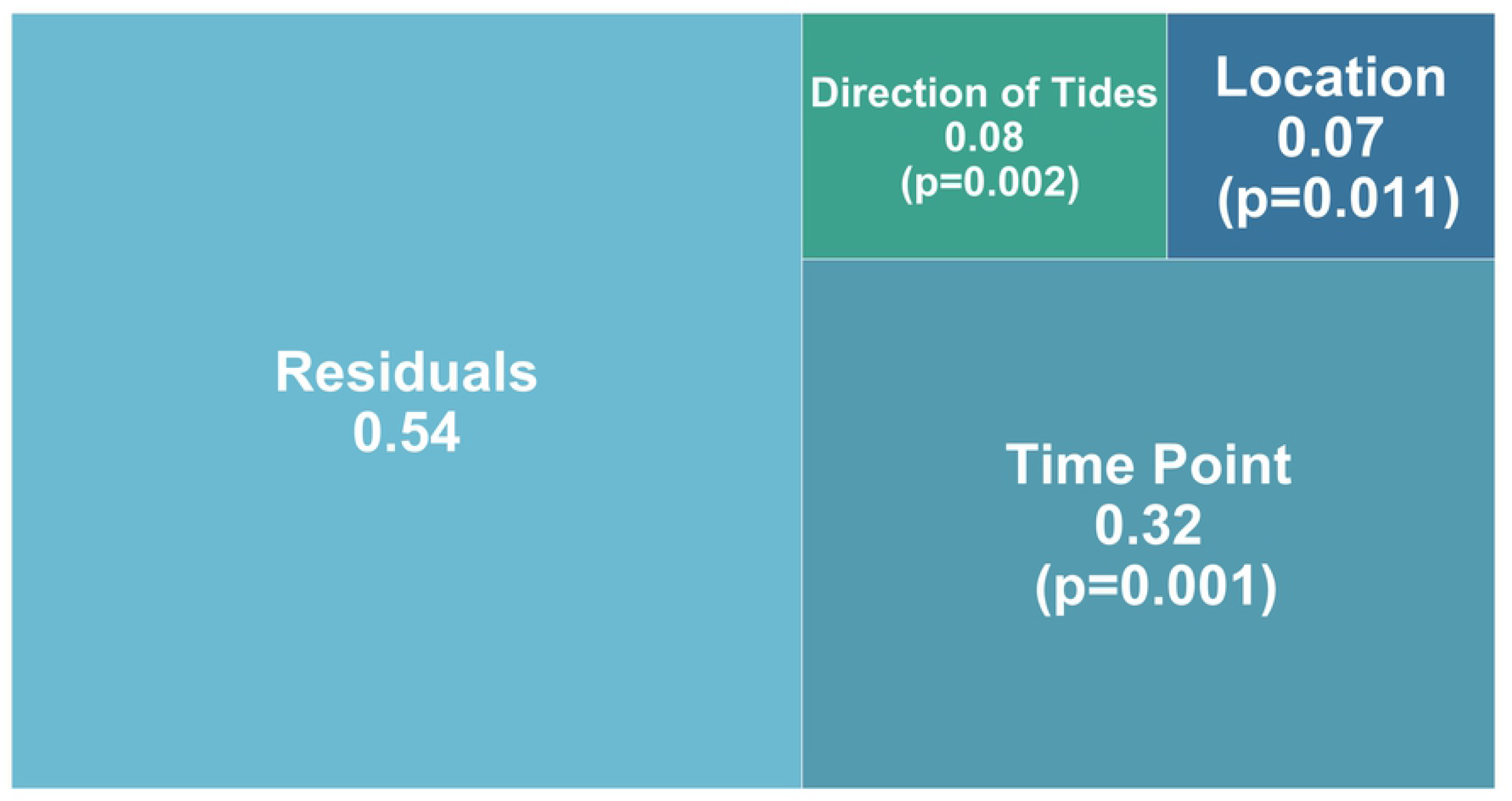
Apportioned variance plot from a PERMANOVA with Bray-Curtis dissimilarities. P-values are stated for each factor. The three processes examined are direction of tide (incoming/outgoing/peak), location (A/B/C), and time point (0-96 hrs).

The mean number of total taxa detected with 3 spatial replicates was 34 taxa [95% CI: 29-39 taxa] and the mean for temporal replicates was 36 taxa [95% CI: 33-38 taxa]. There were no significant differences in total taxa richness captured by spatial and temporal replicates. The mean number of monitored species detected with only 3 spatial replicates was 22 species [95% CI: 19-26 species] and the mean using 3 temporal replicates was 23 species [95% CI: 21-25 species]. As with total species richness, monitored species richness had no significant differences between spatial and temporal replicates.

Species accumulation curves showed that species richness approached saturation, with a species capture estimate of 94.6%, across all 36 samples (Fig 4). Species capture increased to 98.7% when focusing only on species monitored by KFM, Reef Check, and PISCO, and to 99.6% when considering only Reef Check monitored species. Using only 3 random replicate samples, the typical eDNA sampling protocol, species capture estimates were 75.9%, 84.2%, and 92.4%, respectively.

**Fig 4.**
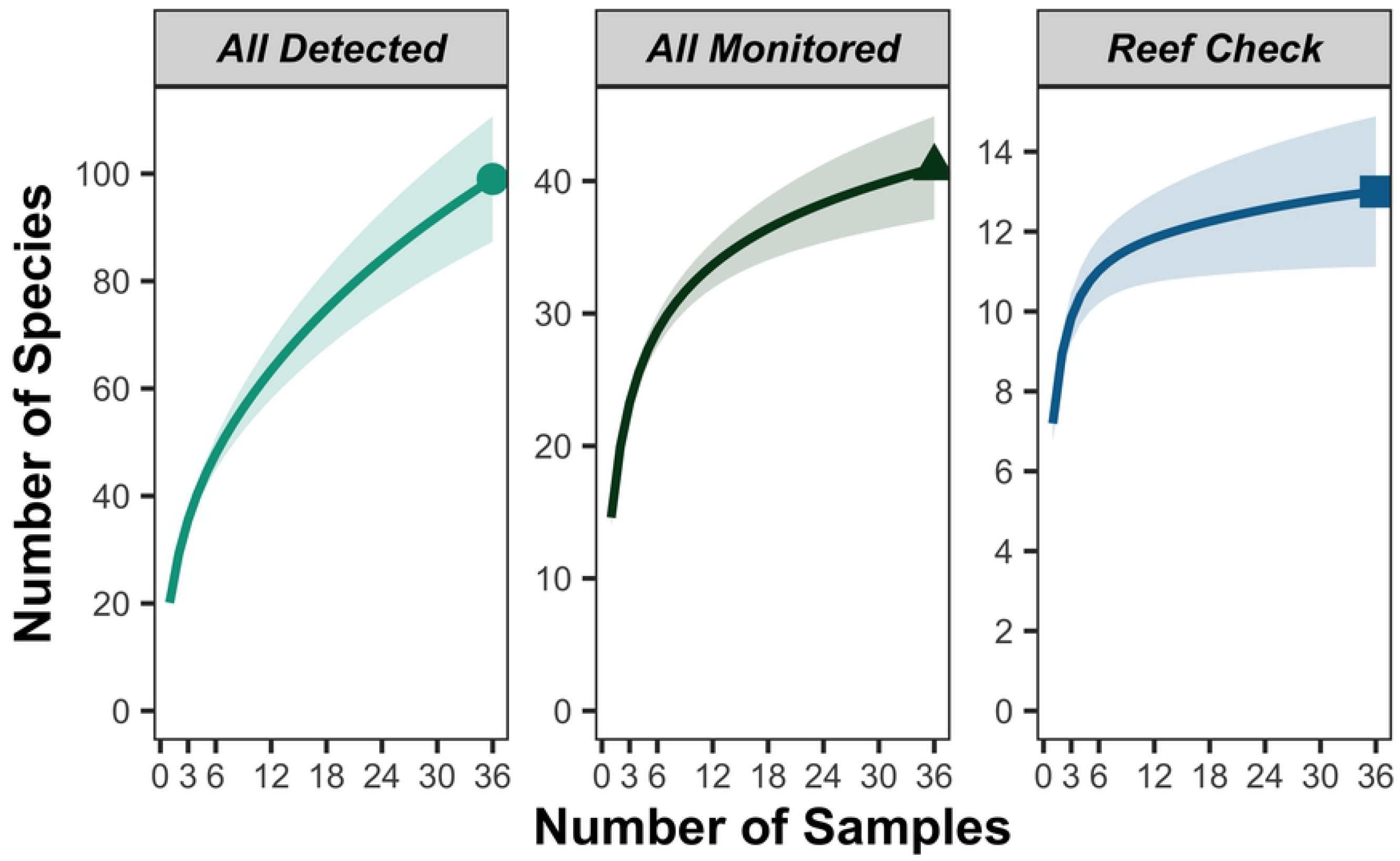
eDNA Metabarcoding Species accumulation curves. Species accumulation curves which indicate how many species on average are detected with increasing number of samples taken. The left graph is a species accumulation curve for total marine vertebrate diversity. This middle graph is a species accumulation curve for fish species that are managed by Reef Check, PISCO, and KFM. The right graph is a species accumulation curve for fish species that are managed by Reef Check.

## Discussion

### Temporal and spatial variation in foreign eDNA signatures

In contrast to studies showing eDNA persisting for multiple days in aquaria and mesocosms [3,8], this study demonstrates that, *in situ*, eDNA signals can be short lived, falling below detection thresholds in only 7.5 hrs. These results are highly similar to results from another temperate region in Japan [11], suggesting that eDNA signals can dissipate rapidly in temperate marine environments through degradation and/or advection. The ephemeral nature of eDNA is further supported by the high variation in the detection of local fish communities sampled repeatedly at the exact same locations over a period of 4 days. Combined, these results suggest that persistence times of eDNA in dynamic marine environments is likely much shorter than currently believed.

While degradation certainly contributed to the dissipation of the introduced eDNA signal over time, advection was also likely a contributing factor. The detection of our introduced eDNA at both sites 19 meters away, only 1.5 hours after release, indicates that the eDNA was moving in both directions at a rate of at least 12 m/hr. While not particularly fast, this rate was fast enough to impact eDNA detection. Previous studies in highly dynamic marine environments show that eDNA can be transported tens of kilometers [40]. However, our study occurred in a protected bay with relatively limited water movement, so advection was not expected to play a major role in eDNA signal detection. This surprising result indicates that eDNA transport may play a significant role in the ability to detect eDNA signals, even in relatively protected marine ecosystems, and may explain the high variability in species detection across time at our fixed sampling locations. Thus, accounting for fine-scale physical oceanography and transport processes will be critical when designing eDNA sampling regimes.

### Temporal and spatial variation in native eDNA signatures

As with the introduced eDNA signature, the detection of the native vertebrate communities was transient and highly variable. eDNA recovered 99 total vertebrate taxa, and the majority of these taxa would be classified as “resident” taxa, such as kelp bass (*P. clathratus*) and garibaldi (*Hypsypops rubicundus*). However, only one species (*C. punctipinnis*) was observed in all samples. A small number of common kelp forest fishes were seen in the majority (>90%) of samples (e.g. *Engraulis mordax, Semicossyphus pulcher, Halichoeres semicinctus*), but the vast majority of taxa were only detected intermittently in eDNA samples (S1 and S2 Figs). Intermittent detection is expected for highly mobile/migratory species observed in our samples such as ocean sunfish (*Mola mola*), California bat rays (*Myliobatis californica*) or Risso’s dolphins (*Grampus griseus*). However, it is surprising that species with demonstrated high site fidelity (e.g. *Oxyjulis californica*) [41] have intermittent eDNA signatures at fixed sampling locations.

Interestingly, time of sampling explained the largest amount of variation in communities recovered by eDNA. The most likely reason for this result is the autocorrelation between time of day and tides over the 4 days of this experiment. However, only 8% of the variation in eDNA signatures was explained by the direction of the tide, indicating that tidal advection was not a major driver of temporal variation in eDNA signatures at this site. Other processes related to time could include fish behavior and activity patterns. For example, more fish movement could lead to more sloughing of eDNA, or predation on fish like anchovy, sardines, or silversides could result in spikes of eDNA as injured fish release blood or other tissues into the water. Alternatively, given that DNA is broken down by UV light, eDNA degradation could be faster during times of high solar irradiance, however there was no obvious temporal pattern observed in the presence/absence of species in this study.

Although the majority of the detected species were resident demersal fish species, there was substantial variation in the community composition recovered across temporal eDNA sampling. The highly localized nature of this eDNA signal suggests that eDNA applications that require robust, reliable, and repeatable data on community composition (e.g. MPA monitoring) will require multiple temporal and/or spatial replicates to detect all species within a given marine environment. These findings are unsurprising given the nature of species distributions in ecosystems and imperfect sampling methods [42].

### Utility of eDNA in detecting monitored taxa

In total, eDNA recovered broader species diversity than visual survey protocols. Across all samples, eDNA detected 85 species, 41 of which are monitored by KFM [43], Reef Check [19], or PISCO [44]. Interestingly, many of these monitored species observed were only found in a single sample (S1 and S2 Figs), indicating stochasticity in eDNA surveys, akin to that of visual surveys. Although we recovered 99 species, this total required 36 samples over 96 hours. Typically, eDNA studies use only 3 biological replicates at a single time point [3,45,46]. When only 3 random samples were used, eDNA captured an average of 36 taxa (Fig 4A) of which an average of 23 were monitored species (Fig 4B). When only using a single, random 1 L sample, eDNA captured an average of 20 taxa of which 15 were monitored species.

Our study design precluded paired visual surveys as diver movement could affect eDNA transport. However, our study site is ~100 m from a site monitored annually by Reef Check through 18 replicate visual surveys on SCUBA along a 30 m long transect [19,47]. Reef Check data from the same year and season as our study reported seeing only 8 of 33 fish species that Reef Check monitors (S4 Table) [19,47] compared to 99 taxa recovered by eDNA. Not only did eDNA recover all eight species reported by Reef Check, but eDNA also detected five more species monitored by Reef Check (*Embiotoca jacksoni, Anistotremus davidsonii, Sphyraena argentea, Paralabrax nebulifer*, and *Labrisomus xanti*) but were all absent in their visual surveys. Importantly, eDNA outperformed visual surveys regardless of number of samples used, averaging 10 Reef Check monitored species in 3 random samples and 7 from a single random eDNA sample (Fig 4C). These results indicate that eDNA consistently recovers more species than visual surveys. However, given the stochasticity of eDNA signals, in limited cases visual surveys may be better, for example detection of specific, less abundant conspicuous indicator taxa that surveyors are trained to look for (e.g. garibaldi).

### Other considerations

Perhaps one of the more perplexing results of this study was the high variation in community diversity observed across space and time in our eDNA samples, and the inability to link these patterns to any physical processes that could explain this variation. Groups overlapped on the Bray-Curtis dissimilarities NMDS ordination plots (S3A and S3B Figs), and there was no clear pattern of prevalence on the heatmaps (Figs 2 and S1). Furthermore, most variation could not be accounted for by any of the three physical oceanographic processes analyzed. Kelly et al. [13] reports that PCR replicates and amplification bias accounted for 12-38% of the variance in intertidal communities, suggesting that PCR variation could be a potential significant driver of our observed variation. Similar results were found by Doi et al. [42] and from simulation results by Kelly et al. [34]. Although we performed PCR in triplicate to limit the impacts of PCR bias, those technical replicates were pooled, rather than being sequenced individually. While we cannot directly assess variation among our PCR amplifications, there is no reason to believe that pooling before indexing (our protocol), versus indexing before pooling (Kelly et al. protocol) should make a difference in terms of variation among sequencing, as ultimately the same amount of product from PCR reactions is loaded onto the sequencer. However, a few studies have shown that three PCR replicates fail to eliminate variation due to PCR bias, with PCR bias still resulting in ≥50% variation [48,49]. Thus although our study design precludes direct observation of variation across PCR replicates and takes measures to reduce PCR bias, PCR bias may still explain a large proportion of the 54% of variation that remains unaccounted for in this study.

Another possibility is that physical movement of fish in this environment could drive the movement of eDNA signatures. Katija and Dabiri [50] showed how diel migration of plankton resulted in biogenic mixing of ocean surface waters, and fish can likely do the same [51]. Providing vertical structure and occurring in a marine protected area, the dock at Wrigley Marine Science Center attracts large numbers of schooling fish (*pers. obs*.). Given how protected from currents, such biogenic mixing could account for the transport of eDNA in multiple directions, and perhaps explain some of the stochasticity in recovered eDNA signals.

An important unintended result of our study is the impact of sampling design on eDNA studies. There is significant variation in eDNA field sampling protocols, with strategies employing different volumes (ranging 1-4 liters; [3,12,14,46]), biological replicates at a single sampling point (ranging 1-4 replicates;[12,46,52]), spatial replicate within an ecosystem (ranging 1-30 replicates; [3,12,14,15]), and combinations of these approaches, all designed to capture a greater proportion of total species diversity. For our study, we collected a single liter of sea water at three points along a transect at 19 m intervals. Community analysis recovered a surprisingly limited number of taxa in any given sample, ranging from 19 to 21 taxa (mean = 20). There was also high variability with the foreign eDNA signature. Despite this variation, subsequent analyses showed that total species richness recovered did not depend on whether replicates were taken over space or time. The lack of a trade-off between the two is important for eDNA sampling designs, indicating that multiple spatial samples can recover the same degree of species richness as sampling over multiple days. Compared to temporal replicates, spatial replicates are often easier to collect and cheaper as expenses such as ship time are only needed for a few hours. Thus our findings suggest that a space for time sampling design can help ensure eDNA remains an easy to deploy and affordable monitoring tool.

Surprisingly, even when we combined all 36 spatial and temporal samples, species accumulation curves suggest that additional sampling is needed to detect all vertebrate biodiversity present. Most eDNA studies sample 1-4 biological replicates taken simultaneously and at most 2 separate time points [3,15,16,52,53]. Based on the results of this study, such sparse sampling efforts may not yield the most complete picture of local diversity. Further tests are required to determine whether species detection could be maximized while maintaining efficiency by pooling multiple water samples or multiple eDNA extractions, allowing broader spatial or temporal sampling without increasing the number of PCR and library preparations that would result in significantly greater lab costs.

Although eDNA was able to capture a broad array of local fish diversity, we note that many taxonomic assignments were made to taxa that are not native to California, but which have closely related taxa in California coastal waters. This result highlights a key limitation of eDNA metabarcoding approaches, namely the need for complete and accurate reference databases for taxonomic assignment [23]. Future voucher barcoding efforts are needed to establish more complete and accurate marine vertebrate reference databases to improve the accuracy and effectiveness of eDNA approaches. Similarly, it is important to archive metabarcoding datasets because bioinformatic pipelines like *Anacapa Toolkit* [23] make it straightforward to rerun legacy datasets, as reference databases become more complete.

## Conclusion

While eDNA holds promise to improve the way that we monitor marine biodiversity, there is much to be learned about the dynamics of eDNA in the natural environment. Diffusion and transport of eDNA, not just degradation, impact our ability to detect taxa within the marine environment, resulting in heterogeneity that may not faithfully reconstruct local communities. However, the impacts of these processes can be minimized by increasing the sampling effort across space or time, allowing diversity estimates to converge on the most complete reconstruction of local communities. Likewise, sampling efforts can be scaled down if monitoring focuses on a smaller subset of common taxa. As marine environments worldwide continue to be impacted by anthropogenic stressors and climate change [54], it will be essential to continue to develop and refine eDNA sampling strategies, allowing this method to achieve its promise as a rapid, reliable, repeatable, and affordable tool for marine ecosystem monitoring [55] in support of marine ecosystem management.

## Acknowledgments

We thank Erick Zerecero and Onny Marwayana for assistance in the field and laboratory, Selena McMillan for providing Reef Check California data, and the University of Southern California Wrigley Marine Science Center for supporting our field studies.

## Supplementary information

**S1 Fig. Heat map of all taxa detected.** The heat map is ordered by time point and faceted by location (in order of location A, B, then C). Darker blue indicates higher eDNA index scores, interpreted as higher relative prevalence. White indicates the taxa was not detected.

**S2 Fig. Detection Rates of Monitored Species.** Histogram of the percentage of samples (N=36) each species was detected in for all species monitored by Reef Check, Partnership for Interdisciplinary Studies of Coastal Oceans (PISCO), or National Park Service (KFM) observed.

**S3 Fig. NMDS ordination of community assemblages. (A)** NMDS ordination plot of community assemblages using all taxa observed with Bray-Curtis dissimilarities. The plot is colored and filled by direction of tide (incoming/outgoing/peak). Shapes also correlate with direction of tide (incoming/outgoing/peak). **(B)** NMDS ordination plot of community assemblages using all taxa observed with Bray-Curtis dissimilarities. The plot is colored and filled by time point (0-96hrs).

**S1 Table. Sample volumes filtered.** (N=36)

**S2 Table. Primers Sequences.** Includes the primer name, target species, primer sequence (5’ to 3’), Illumina Nextera index adapter, and target fragment length. Underlined and bolded sequences represent the original MiFish-U primer set.

**S3 Table. Touchdown PCR thermal profile.**

**S4 Table. Decontaminated Data Table**. Each taxa states if it is a known native, if it is monitored by Reef Check, Partnership for Interdisciplinary Studies of Coastal Oceans (PISCO), or National Park Service (KFM), and if it was found in the recent survey by Reef Check at a site nearby the study site. Blank cells indicate “No”.

